# A functional genetic toolbox for human tissue-derived organoids

**DOI:** 10.1101/2020.05.04.076067

**Authors:** Dawei Sun, Lewis Evans, Kyungtae Lim, Emma L. Rawlins

## Abstract

Human organoid systems recapitulate key features of organs offering platforms for modelling developmental biology and disease. Tissue-derived organoids have been widely used to study the impact of extrinsic niche factors on stem cells. However, they are rarely used to study endogenous gene function due to the lack of efficient gene manipulation tools. We have systematically developed and optimised a complete genetic toolbox for tissue-derived organoids. This includes “Organoid Easytag”, our efficient workflow for targeting all types of gene loci through CRISPR-mediated homologous recombination followed by flow cytometry for enriching correctly-targeted cells. Our toolbox also incorporates conditional gene knock-down, or overexpression, using tightly-inducible CRISPR interference and CRISPR activation; the first efficient application of these techniques to tissue-derived organoids. These tools will facilitate gene perturbation studies in tissue-derived organoids providing a functional counterpart to many on-going descriptive studies, such as the Human Cell Atlas Project.

## Main

CRISPR and its related techniques (CRISPR interference and CRISPR activation, abbreviated as CRISPRi and CRISPRa) have transformed the study of gene function in model systems. They have been rapidly adopted in cancer and pluripotent stem cell (PSC) lines (Bowden et al., 2020; Gilbert et al., 2013; Tian et al., 2019), but not in tissue-derived human organoids. We have optimised these genetic tools for use in organoids using a tissue-derived human foetal lung organoid system (Nikolić et al., 2017).

First, we aimed to establish a robust workflow for gene targeting in organoids to facilitate reporter and direct knockout (KO) generation. A recent application of non-homologous end joining (NHEJ) to improve gene targeting in organoids has been successful (Artegiani et al., 2020). However, we adopted a homology directed repair (HDR) strategy as a complementary approach. We reasoned that the recombination-based method allows our strategy to deliver precise genetic manipulation with flexible targeting sites and minimal additional genetic changes. We chose fluorescence as a selection marker, allowing targeted cells to be easily isolated using flow cytometry and chimeric colonies to be identified and removed using a fluorescent microscope.

In order to achieve efficient gene targeting, we first sought to maximise: (1) the efficiency of DNA delivery into organoid cells; (2) the efficiency of site-specific DNA cleavage by the Cas9-gRNA complex. Nucleofection achieved up to 70% transfection efficiency and showed consistency across different organoid lines (Supplementary Fig. 1a, b). To optimize site-specific DNA cleavage, we used nucleofection to introduce the Cas9-gRNA complex into cells in different forms (Supplementary Fig. 1c). Consistent with previous reports Cas9 RNPs out-performed plasmid based Cas9 approaches (Kim et al., 2014; Lin et al., 2014), both in the T7 endonuclease assay and an online CRISPR editing analysis tool (Supplementary Fig. 1c,d). Thus, we adopted nucleofection and ssRNP for downstream experiments. This strategy has the advantage that the RNP is rapidly degraded and should produce minimal off-target effects.

To establish our gene targeting workflow, we first focused on generating an *ACTB*-fusion protein, taking advantage of the abundance of ACTB protein in human foetal lung organoids and a previously-published targeting strategy (Roberts et al., 2017). We designed a repair template to generate an N terminal monomeric (m)*EGFP-ACTB* fusion (Fig. 1a). We set the following rules for repair template design to facilitate efficient and consistent gene targeting: (1) protospacer adjacent motif (PAM) sequence mutated to prevent editing by ssRNP (Paquet et al., 2016); (2) 700 nt to 1000 nt length of each homologous arm (Yao et al., 2018); (3) minimal plasmid size to maximise delivery into organoid cells; (4) monomeric forms of fluorescent protein to avoid undesirable fusion protein aggregates. As expected, 72 hours after nucleofection of the ssRNP and repair template, mEGFP^+^ organoid cells could be enriched by flow cytometry (Fig. 1b, c). These cells were collected and pooled together, but seeded sparsely, and successfully expanded into organoid colonies (Fig. 1d). The mEGFP-ACTB fusion protein localized to cell–cell junctions as expected (Roberts et al., 2017). These small colonies could be further expanded into new organoid lines and 59% of lines (n = 17/29 lines, from N = 2 parental organoid lines, Supplementary Fig. 5g) were correctly targeted. Targeted organoids continued to express the multipotent lung progenitor marker, SOX9 (Fig. 1e). We sought to further increase targeting efficiency using drugs previously reported to enhance HDR (Maruyama et al., 2015; Song et al., 2016; Yu et al., 2015). However, using flow cytometry as a simple assay, none of the drugs tested increased the rate of gene targeting (Supplementary Fig. 2).

**Figure 1.**
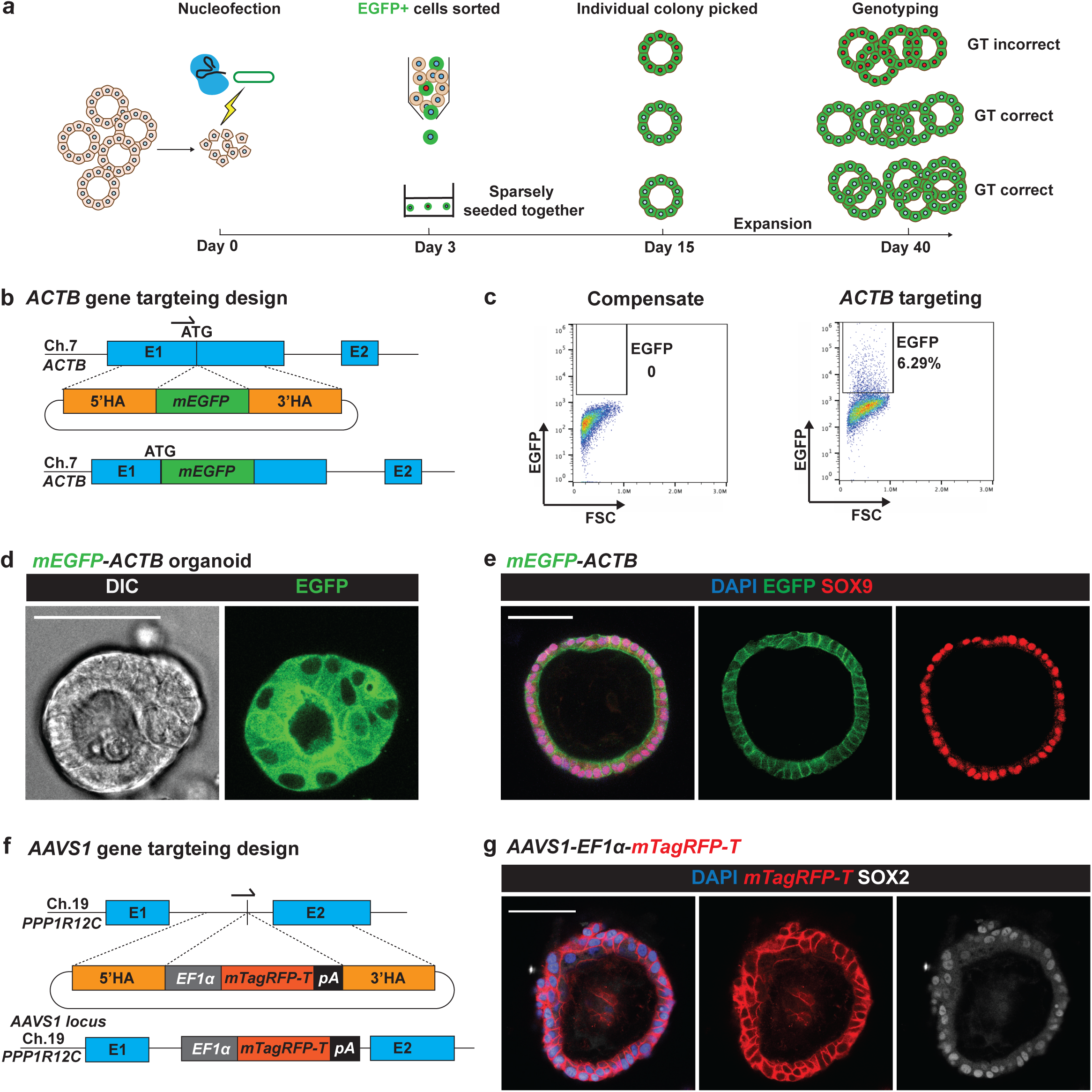
The Organoid Easytag workflow for gene targeting in human organoids. **(a)** Schematic of the Organoid Easytag workflow. ssRNP and a circular plasmid repair template are nucleofected into dissociated cells at day 0. By day 3 cells have proliferated to become tiny colonies and are removed from the Matrigel and dissociated for selection by flow cytometry. EGFP^+^ cells are re-plated sparsely (approx. 1000-1500 cells/well of a 24-well plate) and grown until day 15 when organoids reach a sufficient size to be manually picked under a fluorescent microscope. Organoids are picked into individual wells and passaged until sufficient cells are obtained for both genotyping and freezing down the line. Cells with red nuclei represent incorrectly targeted cells. Cells with white nuclei denote correctly targeted cells. **(b)** Schematic of repair template design for N terminal fusion *mEGFP-ACTB* gene targeting and the final product. Arrow shows the position of gRNA. E1, exon 1; E2, exon 2; 5’HA, 5’ homology arm; 3’HA, 3’ homology arm. **(c)** Representative flow cytometry results showing the percentage of EGFP cells 72 hrs after nucleofection is performed. **(d)** Representative image showing *mEGFP-ACTB* organoid. DIC channel on the left and EGFP channel on the right. **(e)** Immunofluorescence of *mEGFP-ACTB* organoids. Blue: DAPI (nuclei); green: EGFP (ACTB fusion protein); red: SOX9 (lung progenitor marker). (**f**) Schematic of the *AAVS1* targeting repair template design and the final product. E1: Exon1. E2: Exon 2. Arrow indicates the position of the gRNA. **(g)** Immunofluorescence of *AAVS1-EF1a-mTagRFP-T* organoids. Blue: DAPI (nuclei); red: mTagRFP-T (membrane localised reporter); white: SOX2 (lung progenitor marker). Scale bars in all panels denote 50 µm.

The *AAVS1* locus has been considered to be a ‘safe harbour locus’ for expressing exogenous genes in a controllable manner in human cells without silencing (Smith et al., 2008). As a further proof of concept, we successfully targeted *AAVS1* to express a membrane tagged TagRFP-T (mTagRFP-T) for cell shape visualisation (Fig. 1f, 1g). Therefore, we have combined Cas9 RNP with single-stranded synthetic gRNA, a simple circular plasmid repair template and a strategy to enrich correctly targeted cells via flow cytometry for gene targeting in human foetal lung organoids. We named this workflow, Organoid Easytag.

To expand our Organoid Easytag pipeline to reporter generation we targeted *SOX9*. SOX9, a transcription factor, is a tip progenitor cell marker for developing lungs (Nikolić et al., 2017) and *SOX9* reporters are useful for monitoring organoid progenitor state. To overcome its low expression level, we used a Histone H2B-EGFP fusion (H2B-EGFP hereafter) to concentrate the EGFP signal in the nucleus (Fig. 2a). A *T2A* sequence, a self-cleavage peptide, was also inserted between *SOX9* and *H2B-EGFP*, to ensure that SOX9 protein was minimally influenced. This strategy allowed us to enrich correctly targeted cells by flow cytometry. Targeted colonies could be expanded and maintained normal SOX2, SOX9 and NKX2-1 expression (Fig.2b; Supplementary Fig. 3d; Supplementary Fig. 5g). Importantly, we noted that although we were only able to generate *SOX9* reporter lines as heterozygotes (Supplementary Fig 3a, b), the gRNA sites in the wildtype alleles were intact (6/6 lines tested, N=3 parental organoid lines) (Supplementary Fig. 3c). This offers the opportunity to retarget the second allele if desired.

**Figure 2.**
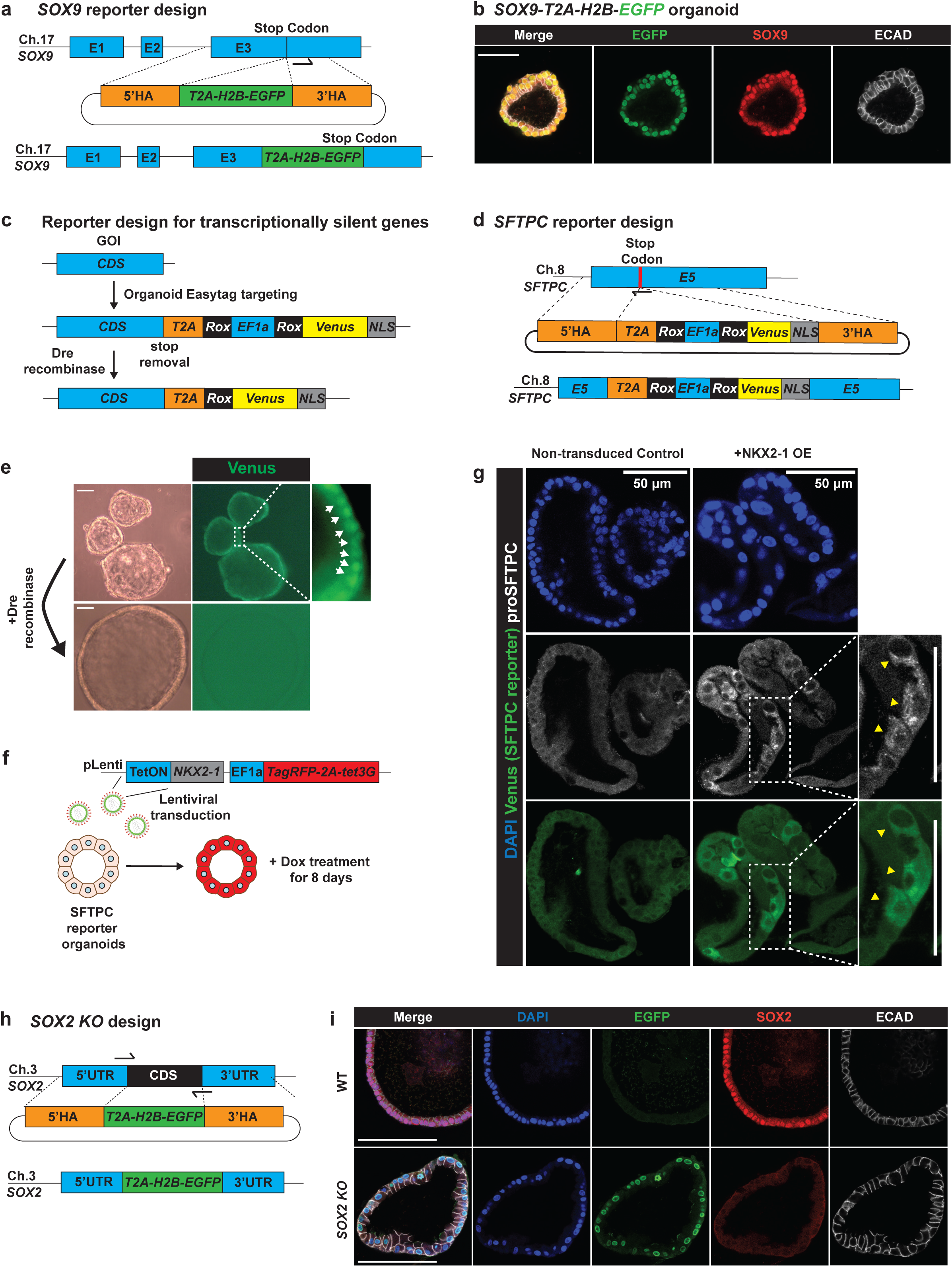
Organoid Easytag workflow facilitates reporter generation and direct knockouts. **(a)** Schematic of the *SOX9* locus repair template design and final product. E1, Exon 1; E2, Exon 2; E3, Exon 3. 5’ HA, 5’ homology arm; 3’ HA, 3’ homology arm. Arrow shows the position of the gRNA. **(b)** Immunofluorescence of *SOX9-T2A-H2B-EGFP* organoids. Green: EGFP (SOX9 reporter); red: SOX9 (lung progenitor marker); white: ECAD (E-cadherin, basal-lateral junctions). **(c)** Strategy for targeting transcriptionally silent genes using the Organoid Easytag workflow. First step: knocking-in to the 3’ end of the locus an exogenous *EF1a* promoter, which is flanked by two Rox sites, driving nuclear-localised Venus expression. A T2A peptide sequence is inserted before the *EF1a* promoter in order to minimise the influence of the future fluorescent reporter on the gene of interest (GOI). Second step: Transient transfection of a plasmid encoding the Dre-recombinase to remove the exogenous *EF1a* promoter. This results in Venus expression becoming dependent on transcription of the targeted locus. (**d**) Schematic of the *SFTPC* locus repair template design and final product. E5, Exon 5; 5’ HA, 5’ homology arm; 3’ HA, 3’ homology arm. Arrow indicates the position of the gRNA. Red bar indicates the position of the stop codon. (**e**) Representative images showing *SFTPC*-*T2A-Rox-EF1a-Rox-Venus-NLS* organoids before and after Dre recombinase expression. Upper panel: nuclear localised Venus fluorescence can be observed under a standard epifluorescent microscope before Dre recombinase expression; bottom panel: Venus fluorescent signal is lost after Dre recombinase expression. Arrows indicate nuclear localised Venus signal. **(f)** Schematic of the strategy for testing *SFTPC* reporter function by NKX2-1 overexpression. *SFTPC* reporter lines were dissociated and transduced with NKX2-1 conditional overexpression lentivirus. Transduced cells are marked by TagRFP expression which can be enriched via flow cytometry. TagRFP expression is coupled to tet3G which, upon doxycycline (Dox) administration, binds to the tetON promoter and drives NKX2-1 overexpression. (**g**) Representative immunofluorescence showing that the *SFTPC* reporter is functional. Blue: DAPI (nuclei); green: Venus (SFTPC reporter); white: proSFTPC protein. (**h**) Schematic showing repair template design and final product for the generation of *SOX2* knockout organoids using Organoid Easytag. Two gRNA sites were used at the N and C terminal of the *SOX2* CDS respectively. *SOX2* CDS was replaced by *T2A-H2B-EGFP*. (**i**) Representative immunofluorescence showing that SOX2 protein is completely knocked-out. Blue: DAPI (nuclei); green: EGFP (*SOX2* transcriptional reporter); red: SOX2 protein (lung progenitor marker); white: ECAD (E-cadherin, basal-lateral junctions). Scale bars denote 50 µm (b, g) and 100 µm (e, i).

We sought to apply Organoid Easytag to transcriptionally silent genes to generate differentiation reporters. We adopted the strategy of inserting an exogenous promoter flanked by two Rox sites which could be subsequently excised by Dre recombinase (Anastassiadis et al., 2009; Roberts et al., 2019). The exogenous *EF1a* promoter drives fluorescent reporter expression for flow cytometry selection allowing us to first positively enrich correctly-targeted cells (Venus^+^), and subsequently enrich (Venus^-^) cells following Dre-mediated *EF1a* removal (Fig. 2c-2e; Supplementary Fig. 4). This design also helps to minimise the repair template size. We targeted the *SFTPC* (surfactant protein C) and *TP63* genes, because they are not expressed in human foetal lung organoid cells and they are well established markers for alveolar type II and basal cell lineages respectively (Barkauskas et al., 2013; Rock et al., 2009). To test *SFTPC* reporter function, we overexpressed NKX2-1 from a lentiviral vector (Fig. 2f) (Attarian et al., 2018). Following NKX2-1 induction, Venus was co-expressed with proSPTPC protein, confirming that the reporter is functional (Fig. 2g). *TP63* reporter organoid lines were produced using the same strategy (Supplementary Fig. 5). These results indicate that the Organoid Easytag workflow can target silent gloci.

The generation of straightforward gene knockouts by induction of indels using the CRISPR-Cas9 system can suffer from translation retention and exon skipping (Smits et al., 2019; Tuladhar et al., 2019). Moreover, in the absence of a strong, immediate phenotype the knockout cells cannot readily be identified. We sought to solve these problems by generating a gene knockout in a controlled manner using Organoid Easytag. We focused on *SOX2* as its function remains to be elucidated in human foetal lung progenitors. We replaced the *SOX2* coding sequence (CDS) with *T2A-H2B-EGFP* to generate *SOX2* knockout organoids. Using two gRNAs targeting the N and C terminal of the *SOX2* CDS respectively, we sequentially replaced both copies of the *SOX2* CDS (Fig. 2h; Supplementary Fig. 6). The EGFP signal was increased after targeting the second copy of the *SOX2* CDS, making selection of serially targeted alleles by flow cytometry straightforward (Supplementary Fig. 6c). *SOX2* knockout colonies proliferate and grow normally, suggesting that SOX2 is not crucial for human foetal lung tip progenitor cell self-renewal (Fig. 2i).

We have demonstrated that our Organoid Easytag pipeline can target various loci, including highly abundant genes, transcription factors, the human safe harbour locus, transcriptionally silent genes and to generate knockouts (summarised in Supplementary Fig. 5g).

Knockouts are not suitable for studying the function of genes which are required for stem cell self-renewal. Moreover, temporal control of gene expression cannot be easily achieved by knockouts. Inducible CRISPRi could potentially solve these problems. We first embedded an N-terminal *KRAB-dCas9* fusion (Mandegar et al., 2016) in a doxycycline (Dox) inducible TetON system aiming to gain temporal control of CRISPRi function. However, the TetON system did not provide tight control of CRISPRi (Supplementary Fig. 7). To solve the leakiness problem we fused KRAB-dCas9 with a destabilising domain derived from *E.coli* dihydrofolate reductase (DHFR) (Fig. 3a). DHFR is stabilised by a small molecule, trimethoprim (TMP) (Fig. 3b) (Iwamoto et al., 2010). We evaluated the new inducible CRISPRi system by depleting a ubiquitous cell surface marker CD71 (Transferrin Receptor C, TFRC). Using previously validated gRNAs (Horlbeck et al., 2016), CD71 protein was depleted in the majority of the cells after 5 days of Dox and TMP treatment (91.4% ± 2.1%, mean ± SEM, N =3, Fig. 3c,d). Whereas no knockdown was observed in the no Dox/TMP treatment group, or in organoids transduced with non-targeting control gRNA. We further evaluated the inducible CRISPRi system using *SOX2*. SOX2 could be efficiently knocked down at both the transcriptional and protein level and no leaky CRISPRi function was observed (Fig. 3e,f), indicating that CRISPRi function has been tightly regulated. Consistent with our *SOX2* knockout (Fig. 2i), we didn’t observe effects on organoid morphology or growth after *SOX2* knockdown (Fig. 3g,h). This further confirmed our finding that SOX2 is not crucial for organoid self-renewal.

**Figure 3.**
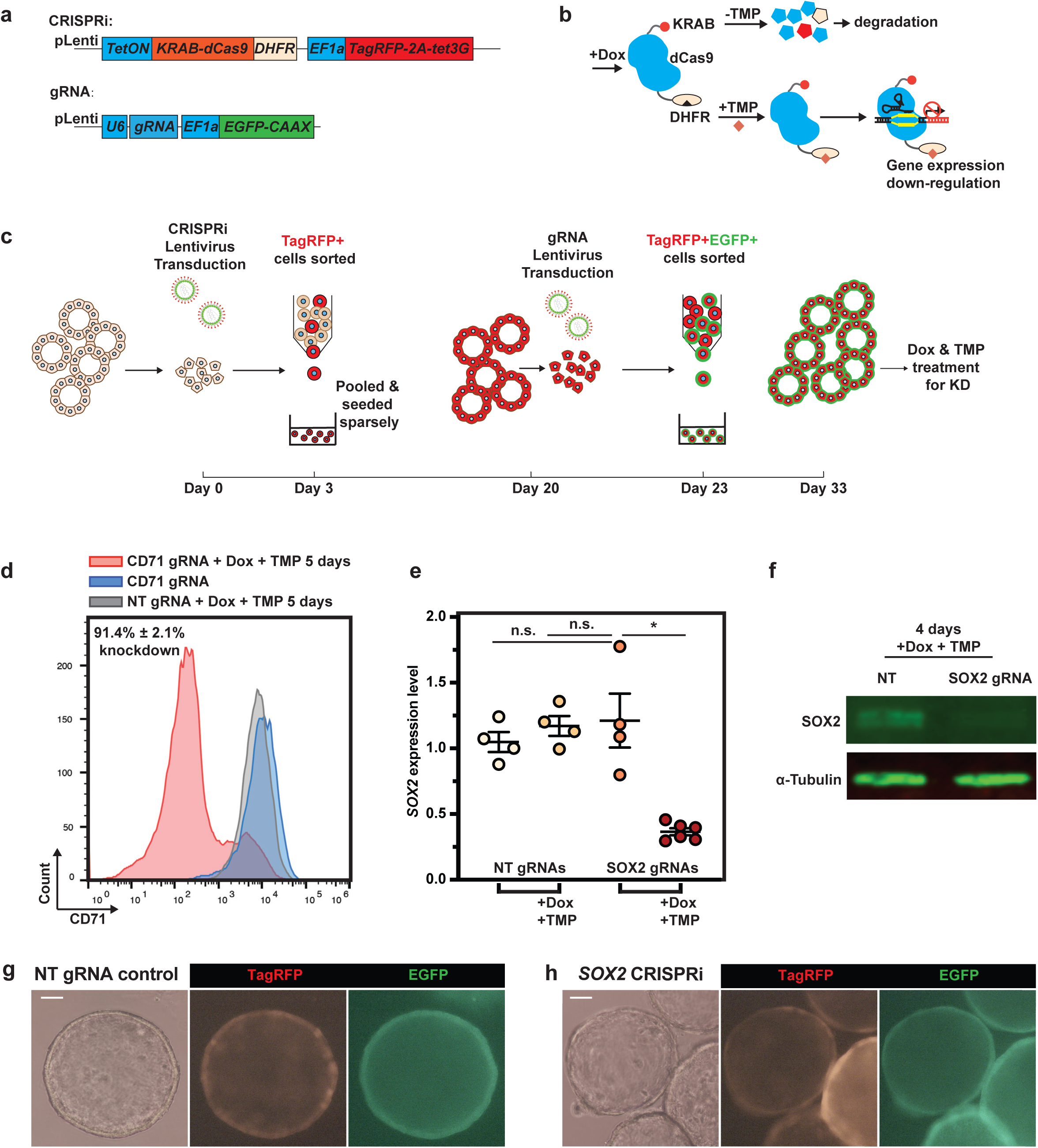
Precise temporal control of CRISPRi in human foetal lung organoids can be achieved via combining the Dox inducible system with a protein destabilising domain. **(a)** Schematic of lentiviral vector designs for sequential introduction of inducible CRISPRi and gRNAs into human foetal lung organoids. N-terminal KRAB-dCas9 was fused with DHFR derived destabilising domain (DD) for stringent control of CRISPRi function. EGFP was fused with a C-terminal CAAX domain to achieve membrane labelling of the transduced cells. **(b)** Schematic showing how the temporal control of CRISPRi protein is achieved by combining the Dox inducible system and DD. The Dox inducible system is somewhat leaky, but in the absence of TMP, any KRAB-dCas9-DHFR fusion protein produced is degraded. Dox treatment results in high levels of KRAB-dCas9-DHFR fusion protein expression. When TMP is not present, the fusion protein is destabilised, whereas when TMP is supplemented, the fusion protein is stabilised and can exert its gene expression repression function. **(c)** Workflow for generating gene knockdown (KD) using CRISPRi system. A parental organoid line with inducible CRISPRi was introduced via lentiviral transduction followed by sorting for TagRFP positive cells. Single cells were re-plated (approx. 3000-5000 cells/well of a 24-well plate) and expanded for around 17 days. gRNA lentivirus was then introduced by a second lentiviral transduction event, followed by sorting for TagRFP/EGFP dual positive cells after 3 days. Cells were re-plated again (approx. 2000-3000 cells/well of a 24-well plate) and expanded for another 10 days before treatment with Dox and TMP. 4-6 days after Dox and TMP treatment gRNA performance was evaluated. **(d)** Representative flow cytometry results for validation of inducible CRISPRi performance using CD71 as target. After TMP and Dox administration for 5 days, CD71 protein was successfully down-regulated. Grey histogram indicates CD71 expression level for organoids with non-targeting control gRNA after Dox and TMP treatment for 5 days; blue histogram indicates the CD71 expression level for organoids with CD71 gRNA without Dox or TMP treatment, showing no KD effect; pink histogram indicates the CD71 expression level for CD71 gRNA with Dox and TMP treatment for 5 days. **(e)** *SOX2* can be effectively knocked-down using inducible CRISPRi without leaky effect indicated by qRT-PCR. Organoids with Dox and TMP treatment were harvested 5 days after the treatment. Each dot represents an independent experiment. 2 independent organoid lines with 2 different non-targeting control gRNAs and 3 different SOX2 gRNAs were used. *SOX2* expression level was normalised to organoids with non-targeting control (NT) gRNAs. Detailed breakdown of these data is shown in Supplementary Figure 8(a). Error bars are plotted to show mean ± SEM. A two-sided Student’s t-test with non-equal variance was used to compare the means of each group. * indicates p-value < 0.05. n.s. indicates non-significant. **(f)** Representative western blot results showing SOX2 was effectively knocked-down using the inducible CRISPRi system. **(g-h)** Representative images showing organoid morphology after *SOX2* knockdown. Images were taken after 5 days of Dox and TMP treatment. *SOX2* knockdown (right panel) didn’t result in obvious phenotypic changes in organoids compared with non-targeting (NT) gRNA control (left panel). Scale bars denote 100 µm.

Finally, we have established the CRISPRa system to switch on endogenous genes in human foetal lung organoids. We tested a previously reported CRISPRa system, dxCas9(3.7)-VPR (Hu et al., 2018), for its ability to activate *SFTPC* transcription (Fig. 4). Inducible CRISPRa can induce *SFTPC* gene expression efficiently at both transcriptional and protein level without influencing organoid growth (Fig. 4b-d).

**Figure 4.**
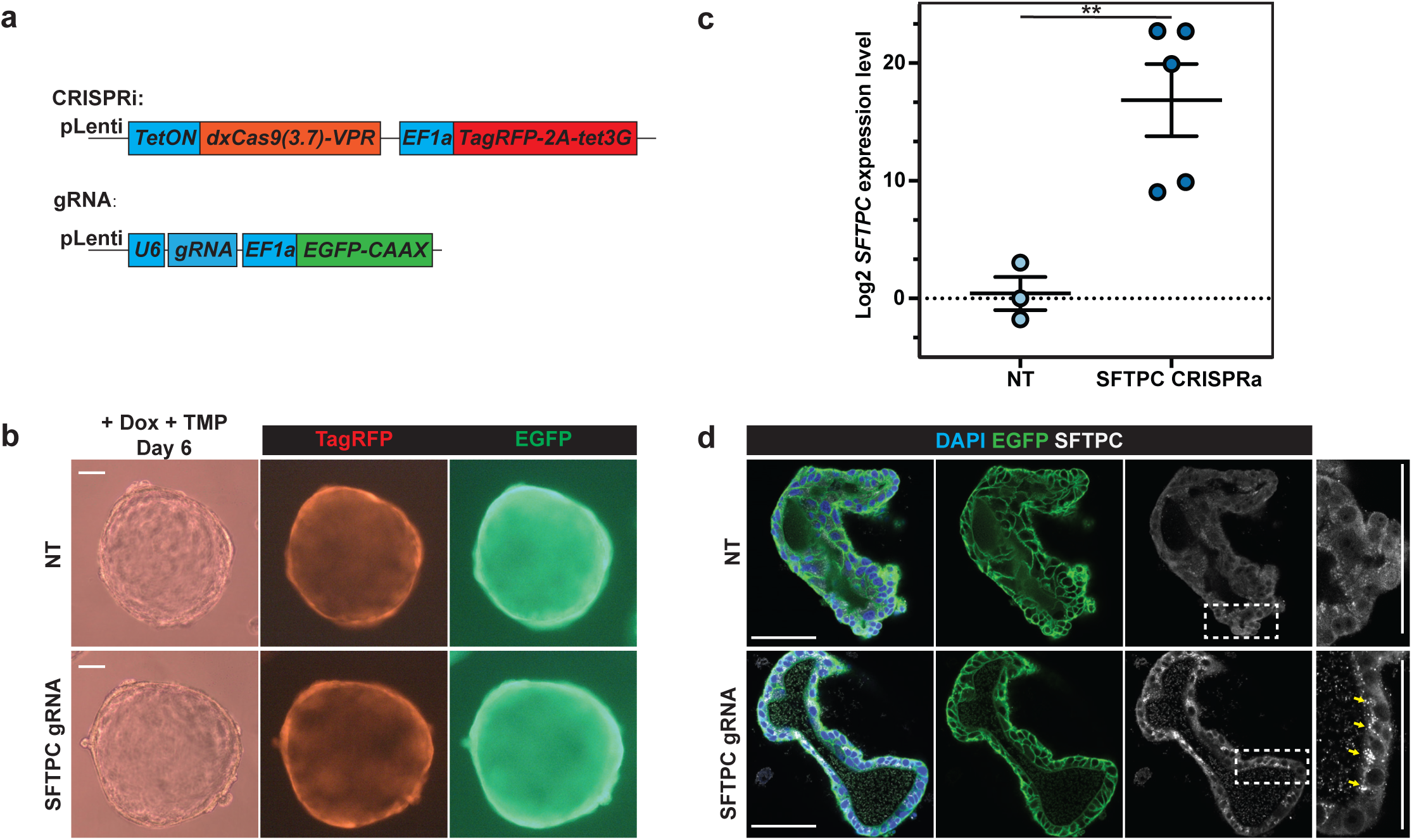
Inducible CRISPRa induced silent gene, *SFTPC*, expression in human foetal lung organoids. **(a)** Schematic of lentiviral design for introducing the CRISPRa system and gRNA into human foetal lung organoids. **(b)** Representative images to show organoid morphology after *SFTPC* activation after 6 days of Dox treatment. *SFTPC* activation (lower panel) didn’t result in obvious phenotypic changes in organoids compared with non-targeting gRNA control (upper panel). **(c)** *SFTPC* can be effectively activated by CRISPRa system indicated by qRT-PCR. Each dot represents an independent experiment. 2 independent organoid lines with 2 different non-targeting control (NT) gRNAs and 3 different *SFTPC* gRNAs were used. *SFTPC* expression level was normalised to organoids with non-targeting control (NT) gRNAs. A detailed breakdown of the results is in Supplementary Figure 8(b). Error bars are plotted to show mean ± SEM. A two-sided Student’s t-test with non-equal variance was used to compare the means of each group. ** indicates p-value < 0.01. **(d)** Representative immunofluorescence showing proSFTPC protein accumulation in organoid cells after CRISPRa activation of *SFTPC* gene expression. Bright SFTPC protein aggregates accumulated in cells, consistent with the hydrophobic nature of SFTPC protein. Green: EGFP. White: proSFTPC protein. Scale bars denote 100 µm (b) and 50 µm (d).

We have established a complete organoid genetic toolbox for gene targeting, reporter generation, controllable gene knockouts, inducible gene knockdown and gene activation in the human foetal lung organoid system. We envision that these tools can be easily adapted for organoid systems derived from other tissues, empowering tissue-derived organoids to benchmark gene function using the Human Cell Atlas as a reference.

## Supporting information

Primer sequences

## Acknowledgements

We thank Dr. Andrew Bassett (Wellcome Sanger Institute) for the Cas9 expressing vector. We thank Prof. Luke Gilbert (UCSF), Prof. Martin Kampmann (UCSF) and Prof. Ruilin Tian (SUSTech) for their advice on CRISPRi and CRISPRa.

D.S. is supported by a Wellcome Trust PhD studentship (109146/Z/15/Z) and the Department of Pathology, University of Cambridge; L.D.E is supported by the Alzheimer’s Research UK Stem Cell Research Centre, funded by the Alborada Trust. K.L. is supported by Basic Science Research Program through the National Research Foundation of Korea (NRF) funded by the Ministry of Education (2018R1A6A3A03012122). E.L.R. is supported by Medical Research Council (MR/P009581/1). Core support from the Wellcome Trust (203144/Z/16/Z) and Cancer Research UK (C6946/A24843).

## Contributions

D.S. designed and performed experiments, analysed data, wrote and edited the manuscript. L.D.E. performed experiments. K.L. performed experiments. E.L.R conceived and led the project, designed experiments, wrote and edited the manuscript.

## Methods

### Derivation and maintenance of human foetal lung organoid culture

Human foetal lung organoids were derived and maintained as previously reported (Nikolić et al., 2017). Briefly, human foetal lung tissues were dissociated using Dispase (8 U/ml Thermo Fisher Scientific, 17105041) at room temperature (RT) for 2 min. Mesenchyme was dissected away using needles. Tips of the branching epithelium were micro-dissected, transferred into 50 µl of Matrigel (356231, Corning) and seeded in one well of a 24 well low-attachment plate (M9312-100EA, Greiner). The plate was incubated at 37°C for 5 min to solidify the Matrigel. 600 µl of self-renewing medium containing: N2 (1: 100), B27 (1: 50), N-acetylcysteine (1.25 mM), EGF (50 ng/ml, PeproTech, AF-100-15), FGF10 (100 ng/ml, PeproTech, 100-26), FGF7 (100 ng/ml, PeproTech, 100-19), Noggin (100 ng/ml, PeproTech, 120-10C), R-spondin (5% v/v, Stem Cell Institute, University of Cambridge), CHIR99021 (3 µM, Stem Cell Institute, University of Cambridge) and SB 431542 (10 µM, bio-techne, 1614), was added. The plate was incubated under standard tissue culture conditions (37°C, 5% CO_2_). Once formed, organoids were maintained in self-renewing medium and passaged by mechanically breaking using P200 pipettes every 10-14 days.

### Whole mount immunostaining for human foetal lung organoids

Organoids were fixed with 4% paraformaldehyde (PFA) directly in the culture plates on ice for 30 min. After two PBS washes, 0.5% (w/v) Bovine Serum Albumin (BSA), 0.2% Triton-X in PBS (washing solution) was added and left on ice overnight to dissolve Matrigel. Organoids were then transferred into multiple CellCarrier-96 Ultra Microplates (PerkinElmer, 6055300) for staining. Subsequently, blocking was performed in 5% donkey serum (Stratech, 017-000-121-JIR), 0.5% (w/v) Bovine Serum Albumin (BSA), 0.2% Triton-X in PBS (blocking solution) at 4°C overnight. For primary antibody staining, the following antibodies in blocking solution were used at 4°C overnight: SOX2 (1: 500, Bio-techne, AF2018), SOX9 (1: 500, Sigma, AB5535), E-cadherin (1: 1500, Thermo Fisher Scientific, 13-1900), NKX2-1 (1: 500, AbCam, ab76013), TagRFP (1: 1000, Evrogen, AB233), GFP (1: 500, AbCam, ab13970), proSPC (1: 500, Merck, AB3786). After washing off the primary antibodies, the following secondary antibodies in washing buffer were used at 4°C overnight: donkey anti-chick Alexa 488 (1: 2000, Jackson Immune, 703-545-155), donkey anti-rabbit Alexa 594 (1: 2000, Thermo Fisher Scientific, A-21207), donkey anti-goat Alexa 594 (1: 2000, Thermo Fisher Scientific, A-11058), donkey anti-goat Alexa 647 (1: 2000, Thermo Fisher Scientific, A-21447), donkey anti-rat Alexa 647 (1: 2000, Jackson Immune, 712-605-153). The following day, DAPI (Sigma, D9542) staining was performed in washing solution at 4°C for 30 min. After two washes with PBS, 97% (v/v) 2’−2’-thio-diethanol (TDE, Sigma, 166782) in PBS was used for mounting. Confocal z stacks were acquired using Leica SP8 at an optical resolution of 1024 × 1024 using a 40x lens. Single plane images are shown. Images were processed using ImageJ (version 2.0.0).

### Plasmid nucleofection

For testing transduction efficiency, organoids were dissociated into single cells using pre-warmed TrypLE™ Express (12605028, Thermo Fisher Scientific) at 37°C for 10 min. The reaction was terminated by adding Advanced DMEM/F12 (12634028, Thermo Fisher Scientific) and cells passed through a 30-micron cell strainer. 2 × 10^5^ organoid single cells were re-suspended with Lonza P3 nucleofection buffer and 1 µl of pmaxGFP (Lonza) and transferred to 20 µl nucleofection cuvette (V4XP-3024, Lonza). Nucleofection was performed using Lonza 4D Nucleofector with X unit using program EA125. After nucleofection, self-renewing medium supplemented with 10 µM Y-27632 (ROCK inhibitor, ROCKi, 688000, Merck) was added to dilute the P3 buffer. Cell mixture was then seeded in Matrigel in 2 wells of a 24-well plate and cultured with self-renewing medium with ROCKi (10 µM) for 72 hrs before FACS analysis.

### Lentiviral production

We grew HEK293T cells in 10-cm dishes to a confluence of 80% before we transfected the lentiviral vector (10 µg) with packaging vectors including pMD2.G (3 µg, Addgene plasmid # 12259), psPAX2 (6 µg, Addgene plasmid # 12260) and pAdVAntage (3 µg, E1711, Promega) using Lipofectamine 2000 Transfection Reagent (11668019, Thermo Fisher Scientific) according to manufacturer’s protocol. After 16 hrs, medium was refreshed. Supernatant containing lentivirus was harvested at 24 hrs and 48 hrs after medium refreshing and pooled together. Supernatant was centrifuged at 300g to remove cell fragments and passed through 0.45 µm filter. Lentivirus containing > 10 kb length insert (inducible CRISPRi and CRISPRa) was concentrated using AVANTI J-30I centrifuge (Beckman Coulter) with JS-24.38 swing rotor at 72000g for two hours at 4 degree. Lentivirus pellets were dissolved in 200 µl PBS. Other Lentivirus, including gRNA and NKX2-1 inducible overexpression, was concentrated using Lenti-X™ Concentrator (631232, Takara). Lentivirus pellets were dissolved in 400 µl PBS.

### Lentivirus infection of organoids to test infection efficiency

Organoid single cell suspension was prepared as for nucleofection. 5 µl lentivirus (CMV-myrAKT-IRES-GFP) suspension was applied to 2 × 10^5^ organoid single cells suspended in 500 µl self-renewing medium with 10 µM ROCKi (without Matrigel) in one well of 24-well plate and incubated at 37 °C overnight. The following day, cells were harvested and washed twice with PBS before pelleting and seeding in Matrigel in two wells of 24-well plate. Cells were grown in self-renewing medium with ROCKi (10 µM) for 72 hrs before flow cytometry. CMV-myrAKT-IRES-GFP was only used for checking lentiviral transduction efficiency (Supplemental Fig. 1).

### Lipofectamine transfection of organoids

Organoid single cells were prepared the same way as for nucleofection. For comparing transduction efficiency, 1 µg of pmaxGFP (Lonza) was mixed with 1 µl of Lipofectamine™ Stem Transfection Reagent (STEM00001, Thermo Fisher Scientific) according to manufacturer’s protocol. 50 µl reaction mixture was applied to 2 × 10^5^ organoid single cells suspended with 450 µl self-renewing medium with ROCKi (without Matrigel) in a single well of a 24-well plate. The plate was then centrifuged at 32°C at 600g for 1 hr, followed by incubation at 37°C for 2-4 hrs. Cells were then harvested, pelleted and seeded in Matrigel in two wells of a 24-well plate and grown in self-renewing medium supplemented with ROCKi (10 µM) for 72 hrs before FACS analysis.

### Nucleofection for gene targeting

Cas9 protein was prepared and used as previously reported (Bruntraeger et al., 2019). If synthetic single strand gRNAs were used, 2 µl spCas9 (4 µg/µl) and 2.88 µl of ssRNA (100 µM, Synthego) were mixed and incubated at RT for a minimum of 10 min in order to form ssRNPs. If synthetic cr/tr RNA heterodimers were used, 200 pmol synthetic cr RNA (IDT) and 200 pmol synthetic tr RNA (IDT) were mixed with 2.5 µl Nuclease Free Duplex Buffer (11-01-03-01, IDT) and denatured at 95 °C for 2 min. 2 µl of cr/tr RNA heterodimer was cooled down to RT on the bench, mixed with 2 µl spCas9 (4 ug/ul) and incubated at RT for a minimum of 10 min to form cr/tr RNPs. At the same time organoids were dissociated into single cells, according to the protocol previously described for nucleofection. 4 × 10^5^cells were suspended using Lonza Nucleofection P3 buffer, mixed with 10 µg of appropriate plasmid repair template, or with 500 pmol ACTB-GFP11 ssODN (Ultramer DNA Oligos, IDT) with 5 µg CMV-GFP(1-10) plasmid. The cell suspension was further mixed with pre-formed Cas9 RNPs and equally distributed into two 20 µl cuvettes (V4XP-3024, Lonza). Nucleofection was performed using program EA104. After nucleofection, self-renewing medium with ROCKi was added to dilute the P3 buffer. Cell mixture was then taken out and seeded in Matrigel in 4 wells of a 24-well plate and cultured with self-renewing medium with 10 µM ROCKi for 72 hrs before flow cytometry.

gRNA sequences used for gene targeting are as follows:

*ACTB 5’-GCTATTCTCGCAGCTCACCA <u>TGG</u>*, *SOX9 5’-CTTGAGGAGGCCTCCCACGA <u>AGG</u>*, *AAVS1 5’-GTCCCCTCCACCCCACAGTG <u>GGG</u>*, *SOX2 N terminal 5’-CGGGCCCGCAGCAAACTTCG <u>GGG</u>*, *SOX2 C terminal 5’-CGGCCCTCACATGTGTGAGA <u>GGG</u>. SFTPC 5’-GCGTCCTAGATGTAGTAGAG <u>CGG</u>. TP63 5’-TGATGCGCTGTTGCTTATTG <u>CGG</u>.*

PAM sequences are underlined.

### Small molecule influence on gene targeting efficiency

mEGFP-ACTB gene targeting was performed as previously described. After nucleofection, DMSO (0.6 ul, D2650, Sigma), RS-1 (10 uM, R9782, Sigma), L755507 (5 uM, SML1362, Sigma), or SCR-7 (100 uM, SML1546, Sigma) were added to self-renewing medium with ROCKi for 48 hrs. Organoid cells were analysed by flow cytometry 72 hrs after nucleofection.

### T7 Endonuclease Assay

To test for site specific DNA cleavage using the T7 endonuclease assay, organoid cells were harvested 48 hrs after nucleofection of ssRNP, tr/cr RNP, plasmid encoding Cas9 and gRNA or WT control organoids. Genomic DNA was extracted using QIAamp Fast DNA Tissue Kit (51404, Qiagen). PCR was performed using PrimeSTAR® GXL DNA Polymerase (R050A, Takara) with 20 ng of genomic DNA as template according to manufacturer’s protocol. Forward primer: 5’-*TTGCCAATGGGGATCGCAG*-3’ and reverse primer: 5’-*GCTCGATGGGGTACTTCAGG*-3’ were used for *ACTB* locus amplification. 10 µl of PCR product was then mixed with 1.5 µl 10X NEBuffer 2 (B7002S, NEB) and 1.5 µl of Nuclease-free water. The mixture was denatured at 95°C for 10 min, followed by ramp −2°C per second from 95°C to 85°C and ramp −0.3 °C per second from 85°C to 25°C. 2 µl T7 Endonuclease I (1 U/ul, M0302S, NEB) was added and incubated at 37°C for 1 hr. 2.5% Agarose gel was used for electrophoresis.

### ICE analysis for Indel production

Genomic DNA was extracted from organoid cells which were harvested 48 hrs after nucleofection of ssRNP, tr/cr RNP, plasmid encoding Cas9 and gRNA or WT control organoids using QIAamp Fast DNA Tissue Kit (51404, Qiagen). PCR was performed using PrimeSTAR® GXL DNA Polymerase (R050A, Takara) with 20 ng of genomic DNA as template according to manufacturer’s protocol. Forward primer: 5’-*TTGCCAATGGGGATCGCAG*-3’ and reverse primer: 5’-*GCTCGATGGGGTACTTCAGG*-3’ were used for *ACTB* locus amplification. PCR products were cleaned up using Macherey-Nagel™ NucleoSpin™ Gel and PCR Clean-up Kit (Macherey-Nagel, 740609.50) and sent for Sanger Sequencing (Department of Biochemistry, University of Cambridge) using reverse primer: 5’-*GCTCGATGGGGTACTTCAGG*-3’. Sanger sequencing results were compared using ICE online CRISPR editing analysis tool: https://www.synthego.com/products/bioinformatics/crispr-analysis

### Flow cytometry analysis

Organoid single cells were prepared 72 hrs after nucleofection, lentivirus transduction or Lipofectamine transfection. Cells were analysed using Sony SH800S Cell Sorter and Flowjo software (version 10.4).

### Organoid Genotyping

Organoids from a single 48-well plate well were used for genomic DNA extraction with QIAamp Fast DNA Tissue Kit (51404, Qiagen) according to manufacturer’s protocol. PCR was performed using PrimeSTAR® GXL DNA Polymerase (R050A, Takara) with 20 ng of genomic DNA as template according to manufacturer’s protocol. Primers used are listed in genotyping primers file. For each gene targeting, 3 randomly picked lines were chosen for further Sanger Sequencing. 5’ and 3’ homologous arms of the gene targeting product were amplified using PrimeSTAR® GXL DNA Polymerase with aforementioned primers. PCR products were cleaned up using Macherey-Nagel™ NucleoSpin™ Gel and PCR Clean-up Kit (Macherey-Nagel, 740609.50) and sequenced using Sanger Sequencing (Department of Biochemistry, University of Cambridge).

### Western blot

Organoids were harvested, washed twice with Advanced DMEM/F12 and then twice with PBS before pelleting at 300g for 5 min. Organoid cell pellets were re-suspended in 100 µl-200 µl of RIPA buffer with protease inhibitor (Thermo Fisher Scientific, 87786) added. Organoid suspension was incubated for 30 minutes on ice, with strong vortex every 5 minutes. Cell pieces and debris were removed by centrifugation at 13000 rpm. Supernatant was harvested. Protein concentration was measured by BCA assay (Thermo Fisher Scientific, 23227). Equal amount of each protein sample was mixed with Sample Buffer (Bio-rad, 1610747) and beta-mercaptoethanol according to manufacturer’s protocol. Mixture was heated at 95 °C for 5 minutes and cooled down to room temperature.

Samples were then separated on a 4%–12% SDS-PAGE and transferred to nitrocellulose membranes. Membrane was blocked by PBS with 5% BSA for 1 hour at room temperature. Proteins were detected by incubation with primary antibodies (SOX2, Bio-techne, AF2018, 1: 1000 and α-Tubulin, Merck, T6199, 1:2000) and secondary antibodies (Donkey anti-Goat IRDye® 800CW, AbCam, ab216775, 1: 1000; Donkey anti-Mouse IRDye® 800CW, ab21677, 1: 1000). Protein bands were visualised using Li-Cor Odyssey system.

### Lentivirus infection of organoids for inducible knockout and activation

Organoids were dissociated into small organoid pieces or single cells using pre-warmed TrypLE™ Express at 37°C for 5-10 min. Organoid cells were then spun down at 300g for 5 min. The cell pellets from one 24-well-plate well were responded in 500µl self-renewing medium with ROCKi. 20ul of inducible CRISPRi or CRISPRa lentivirus was added and mixed well. The mixture was incubated at 37 degree overnight. The next morning, organoid cells were pelleted at 300g for 5 min, washed twice with PBS and seeded in 100µl Matrigel into 2 24-well-plate wells. Organoid cells were cultured with self-renewing medium with ROCKi for 3 days before dissociated with TrypLE™ Express into single cells for cell sorting. TagRFP^+^ cells were sorted using Sony SH800S Cell Sorter and pooled together and seeded in Matrigel at approx. 3000-5000 cells/well of a 24-well plate. Organoids cells were then expanded around 17 days using self-renewing medium with ROCKi. At this stage, organoids with inducible CRISPRi and CRISPRa system could be frozen as parental lines.

Organoids with inducible CRISPRi and CRISPRa system were broken into small organoid pieces or single cells similarly and transduced with gRNA lentivirus (5µl-10µl virus/500µl organoid cell resuspension). The mixture was incubated overnight and the next morning, organoid cells were pelleted, washed and seeded with Matrigel as described above. After 3 days of culturing with self-renewing medium supplemented with ROCKi, TagRFP and EGFP double positive cells were sorted and seeded in Matrigel at approx. approx. 2000-3000 cells/well of a 24-well plate and expanded for 10-17 days before turning on inducible CRISPRa and CRISPRi. Doxycycline (2 µg/ml, Merck, D9891) and trimethoprim (10 nmol/L, Merck, 92131) were supplemented in self-renewing medium accordingly. Medium was refreshed every 48 hours.

For evaluation of inducible CRISPRi using *CD71* as target, single cells were prepared as described above after 5 days of Dox and TMP treatment. Cells were pelleted at 300g for 5min and re-suspended in 100 µl PBS with 0.5%BSA and 2mM EDTA. 2.5 µl of PE/Cy7 anti-human CD71 antibody (BioLegend, 334111) was added and incubated at 4 degree for 30 min. Cells were then washed with PBS with 0.5%BSA and 2mM EDTA twice and re-suspended in 300 µl of PBS with 0.5%BSA and 2mM EDTA for flow cytometry analysis.

### RNA extraction, cDNA synthesis and qRT-PCR

Organoids were harvested and washed twice with Advanced DMEM/F12, before 350 µl of RLT buffer was added to lyse organoids. RNA extraction was performed according to manufacturer’s protocol using RNeasy Mini Kit (Qiagen, 74104) with RNase-Free DNase Set (Qiagen, 79254). RNA concentrations were measured by Nanodrop (Thermo Fisher Scientific). cDNA was synthesized with MultiScribe™ Reverse Transcriptase (Thermo Fisher Scientific, 4311235) according to manufacturer’s protocol. cDNA was diluted 1: 25 and 6 µl was used for one qPCR reaction with PowerUp™ SYBR™ Green Master Mix (Thermo Fisher Scientific, A25776). Relative gene expression was calculated using the ΔΔCT method relative to *ACTB* control.

### Plasmid Construction

eSpCas9(1.1) was a gift from Feng Zhang (Addgene plasmid # 71814). ACTB gRNA sequence 5’-GCTATTCTCGCAGCTCACC-3’ was cloned into the vector using BbsI sites. ACTB repair template AICSDP-15: ACTB-mEGFP was a gift from The Allen Institute for Cell Science (Addgene plasmid # 87425). SOX9 repair template was created by Infusion (638909, Takara) cloning to insert SOX9 5’ and 3’ homologous arms in EasyFusion T2A-H2B-GFP plasmid (a gift from Janet Rossant, Addgene plasmid # 112851). AAVS1 repair template was created by Infusion cloning to swap the CAG promoter and Puromycin resistance cassette in plasmid AICSDP-42: AAVS1-mTagRFPT-CAAX (a gift from The Allen Institute for Cell Science, Addgene plasmid # 107580). SOX2 knockout repair template was created by Infusion cloning to insert SOX2 5’ and 3’ homologous arms in EasyFusion T2A-H2B-GFP (a gift from Janet Rossant, Addgene plasmid # 112851). SFTPC targeting repair template was created by Infusion assembly of SFTPC 5’ and 3’ homologous arms together with T2A-Rox-EF1a-Rox-Venus-NLS. TP63 targeting repair template was created by Infusion assembly of TP63 5’ and 3’ homologous arms together with T2A-Rox-EF1a-Rox-Venus-NLS. CMV-Dre-T2A-TagRFP vector was created by Infusion assembly of Dre (a gift from Azim Surani Group, Gurdon Institute, Cambridge) and T2A-TagRFP sequences together. NKX2-1 overexpression vector was created by inserting EF1a-TagRFP-2A-tet3G and tetON-NKX2-1 CDS in 2 steps cloning using Infusion cloning into a pHAGE backbone. The minimal CMV-GFP(1-10) plasmid was created by Infusion cloning of CMV-GFP(1-10) from pcDNA3.1-GFP(1-10) (a gift from Bo Huang (Addgene plasmid # 70219) into a pUC57 backbone. For testing lentiviral transduction efficiency, CMV-myrAKT-IRES-GFP vector was created by Infusion cloning to insert myrAKT from pCCL-Akt1 (a gift from Bi-Sen Ding, Icahn School of Medicine, Mount Sinai) and IRES sequence into pFP945 (a gift from Frederick Livesey, University College London). The Dox inducible CRISPRi vector was created by sub-cloning N-terminal KRAB-dCas9 (a gift from Bruce Conklin, Addgene plasmid # 73498) into the NKX2-1 overexpressing vector using XhoI and BamHI sites. Dox inducible CRISPRi with DD control vector was created by In-fusion cloning of a C-terminal DHFR sequence into the Dox inducible CRISPRi vector using BamHI site. The inducible CRISPRa vector was created by sub-cloning dxCas9(3.7)-VPR (a gift from David Liu, Addgene plasmid # 108383) into the NKX2-1 overexpressing vector using XhoI and BamHI sites. The gRNA entry vector was cloned by infusion cloning of a EF1a promoter into pKLV2-U6gRNA5(BbsI)-PGKpuro2ABFP-W vector (a gift from Kosuke Yusa, Addgene plasmid # 67974) using BamHI and EcoRI sites, and then cloned a EGFP-CAAX to swap the EGFP sequence using XhoI and NotI sites. All plasmids will be deposited in Addgene.

---

**Supplementary Figure 1.**
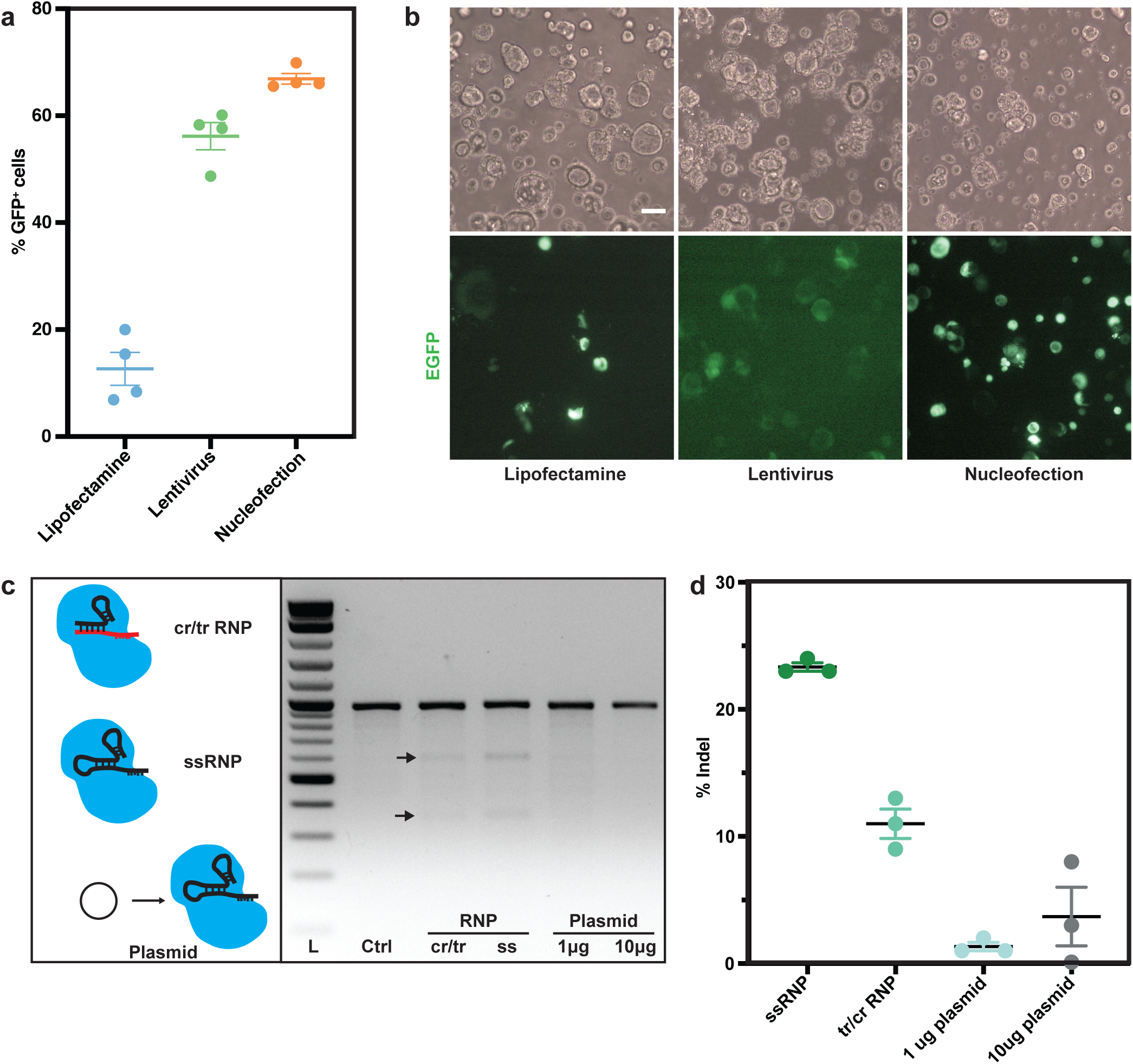
Optimisation of DNA delivery and CRISPR cutting. **(a)** Efficiency quantification of different DNA delivery methods. EGFP positive cells were quantified by flow cytometry 72 hrs after transfection or transduction. N = 4 different organoid lines were used for each condition. Error bars are plotted to show mean ± SEM. **(b)** Representative images showing DNA delivery efficiencies of different methods 72 hrs after transfection/transduction. Top panel: wide-field images; bottom panel: GFP channel. Scale bar denotes 100 µm. **(c)** T7 endonuclease assay showing the DNA cleavage efficiency of different CRISPR methods on the *ACTB* locus. Arrows denote lower bands generated by T7 endonuclease cutting. Left panel: schematic showing the different methods tested for introducing the Cas9 complex. cr/tr RNP, synthetic crispr/tracer RNA heterodimer with Cas9 RNP. ssRNA, single strand synthetic guide RNA with Cas9 RNP. **(d)** Quantification of indels produced by the different CRISPR methods tested using ICE online analysis software (https://ice.synthego.com/). N =3 different organoid lines were used for each condition. Error bars are plotted to show mean ± SEM.

**Supplementary Figure 2.**
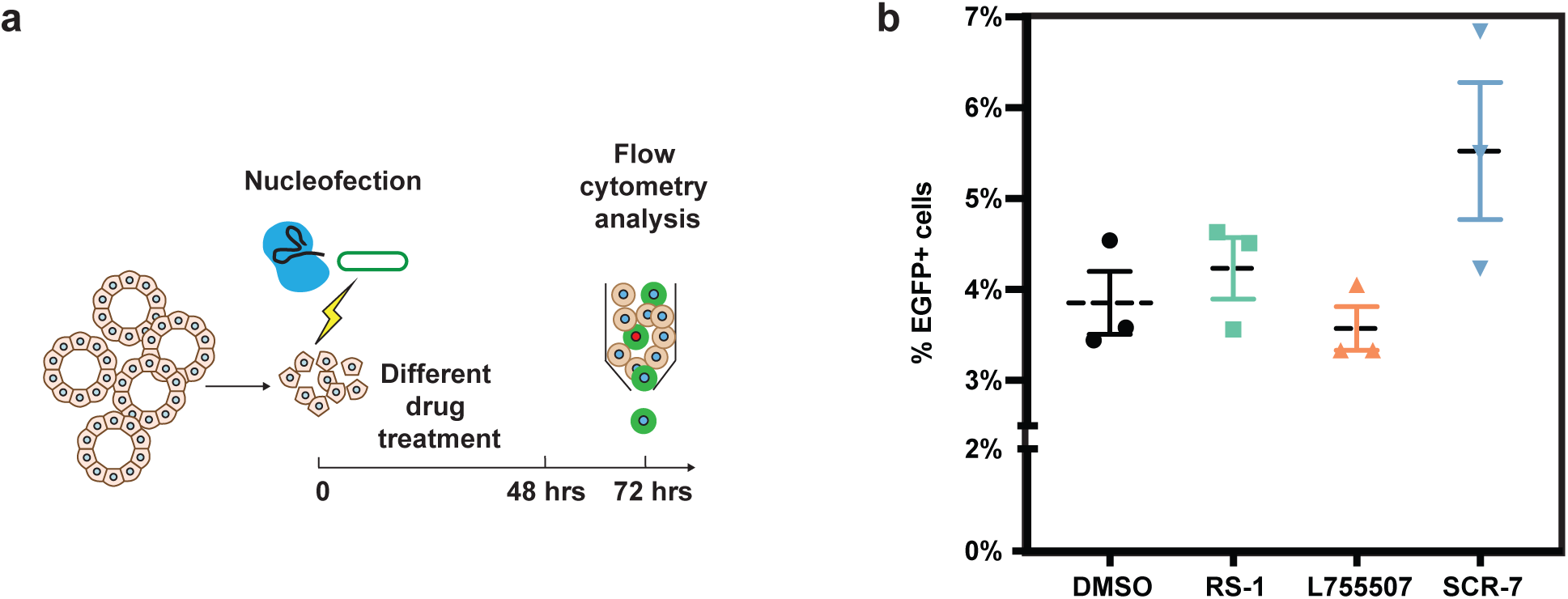
Small molecules did not significantly improve organoid gene targeting efficiency. **(a)** Workflow for testing different HDR enhancing drugs. Organoid cells were treated with DMSO, RS-1, L755507 and SCR-7 for 48hrs after nucleofection for *mEGFP-ACTB* gene targeting. The percentage of EGFP positive cells was analyzed 72 hrs after nucleofection. DMSO treatment was used as a negative control. (**b**) Summary of percentage of EGFP positive. No significant improvement of targeting efficiency was observed after the drug treatments. N =3 different organoid lines were used. Error bars are plotted to show mean ± SEM.

**Supplementary Figure 3.**
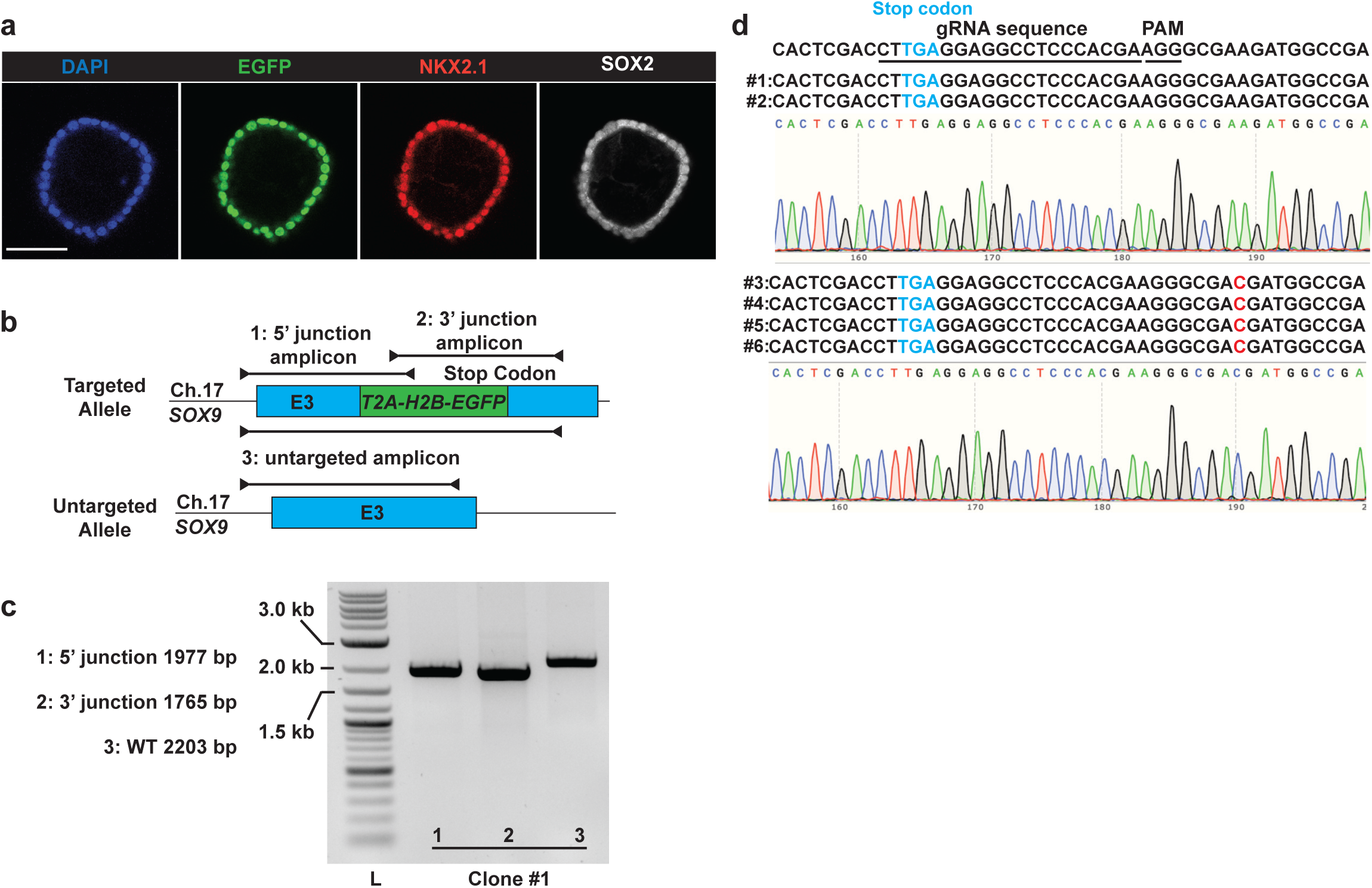
Characterisation of *SOX9* targeted colonies. **(a)** Immunofluorescence of a *SOX9-T2A-H2B-EGFP* clone. Blue: DAPI (nuclei); green: H2B-EGFP (*SOX9* transcriptional reporter); red: NKX2-1 (lung epithelial marker); white: SOX2 (lung progenitor marker). Scale bar denotes 50 µm. **(b)** Schematic of the *SOX9* genotyping strategy. 5’ and 3’ junction amplicons consist of a primer inside the EGFP sequence and another primer upstream, or downstream, of the homology arm respectively. **(c)** Representative gel image showing correct colony genotyping results. **(d)** Illustration of Sanger sequencing results for *SOX9-T2A-H2B-EGFP* heterozygous WT allele, focusing on the gRNA cutting sites. A single point mutation outside the gRNA sequence in the 3’UTR of SOX9 is observed in 4/6 of the clones sequenced (#3-#6). Results were from 6 colonies, randomly picked from N = 3 different organoid lines.

**Supplementary Figure 4.**
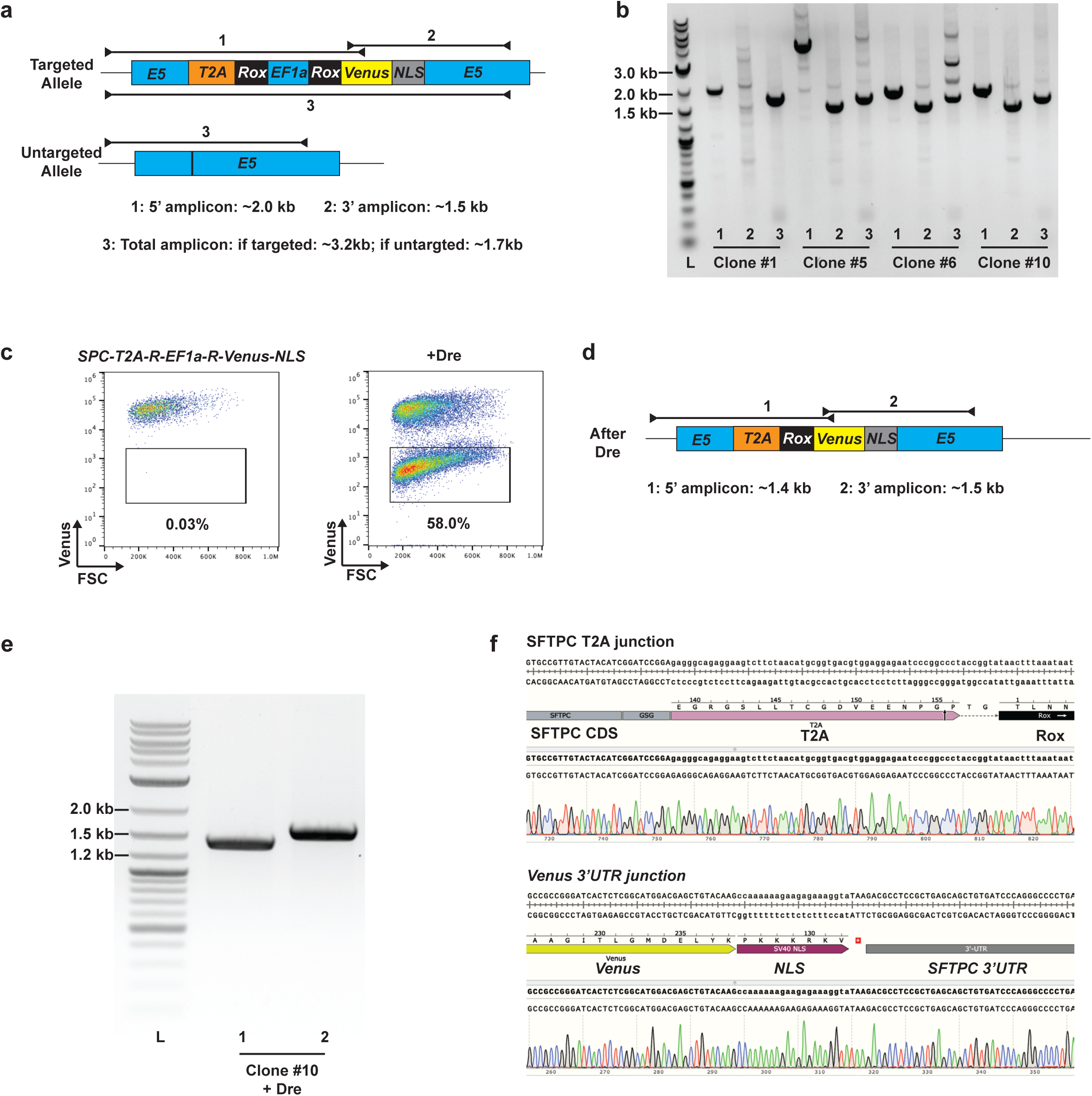
Organoid Easytag to target the *SFTPC* locus. **(a)** Schematic of the *SFTPC* reporter genotyping strategy. 5’ and 3’ junction amplicons consist of a primer inside the Venus sequence and another primer upstream, or downstream, of the homology arm respectively. (**b**) Representative gel image showing correct and incorrect colony genotyping results. Clone #10 was correctly targeted; Clone #6 appeared correctly targeted in gel, however, Sanger sequencing revealed a deletion in the T2A sequence. Clone #1 and #5 were incorrectly targeted. **(c)** Representative flow cytometry results showing loss of the Venus signal after transient expression of Dre recombinase to excise the *EF1a* promoter. **(d)** Schematic of the final *SFTPC* reporter genotyping strategy to validate *EF1a* removal. 5’ and 3’ junction amplicons consist of a primer inside the Venus sequence and another primer upstream, or downstream, of the homology arm respectively. **(e)** Representative gel image showing correct SFTPC reporter colony (after Dre recombinase) genotyping results. **(f)** Sanger Sequencing traces showing the correct *SFTPC-CDS-T2A* and *Venus-SFTPC* 3’UTR junctions.

**Supplementary Figure 5.**
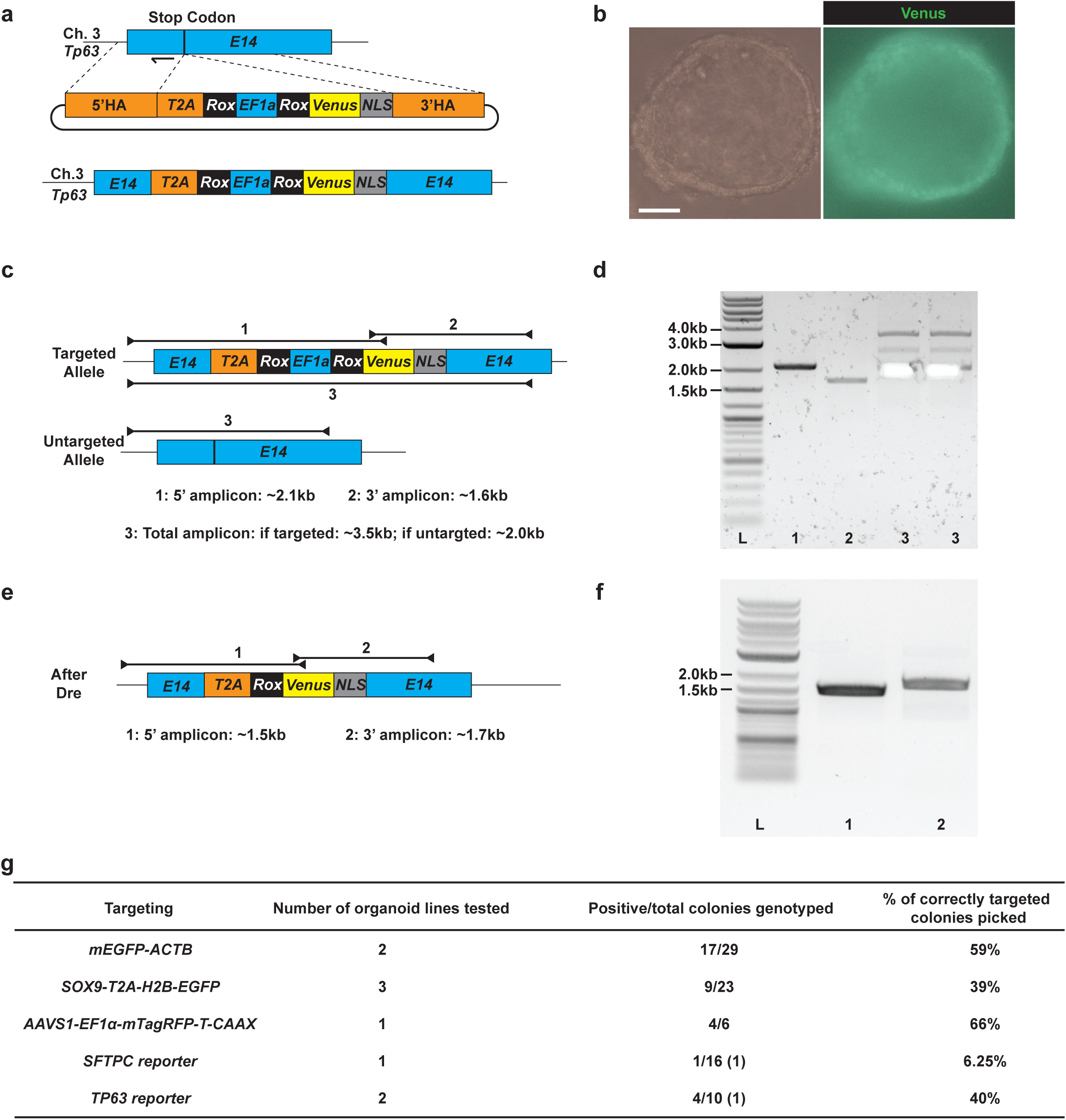
Organoid Easytag to target the *TP63* locus. **(a)** Schematic of the *TP63* locus repair template design and final product. E14, Exon 14; 5’ HA, 5’ homology arm; 3’ HA, 3’ homology arm. Arrow indicates the position of the gRNA. **(b)** Representative images showing *TP63-T2A-Rox-EF1a-Rox-Venus-NLS* heterozygous organoid. Nuclear localised Venus fluorescence can be observed under epifluorescent microscope. Scale bar indicates 50 µm. **(c)** Schematic of the *TP63* genotyping strategy. 5’ and 3’ junction amplicons consist of a primer inside the Venus sequence and another primer upstream, or downstream, of the homology arm respectively. The expected lengths of each amplicon are labelled below the schematic. **(d)** Representative gel image showing correct colony genotyping results. The WT bands were excised for Sanger sequencing in lanes #3. **(e)** Schematic of the final *TP63* reporter genotyping strategy to validate *EF1a* removal. **(f)** Representative gel image showing correct colony genotyping results after Dre recombinase. (**g**) Summary of all the gene targeting results. For *SFTPC* and *TP63* reporter generation, results for the first step gene targeting were shown. The number of lines that were generated after Dre recombinase expression is shown in brackets.

**Supplementary Figure 6.**
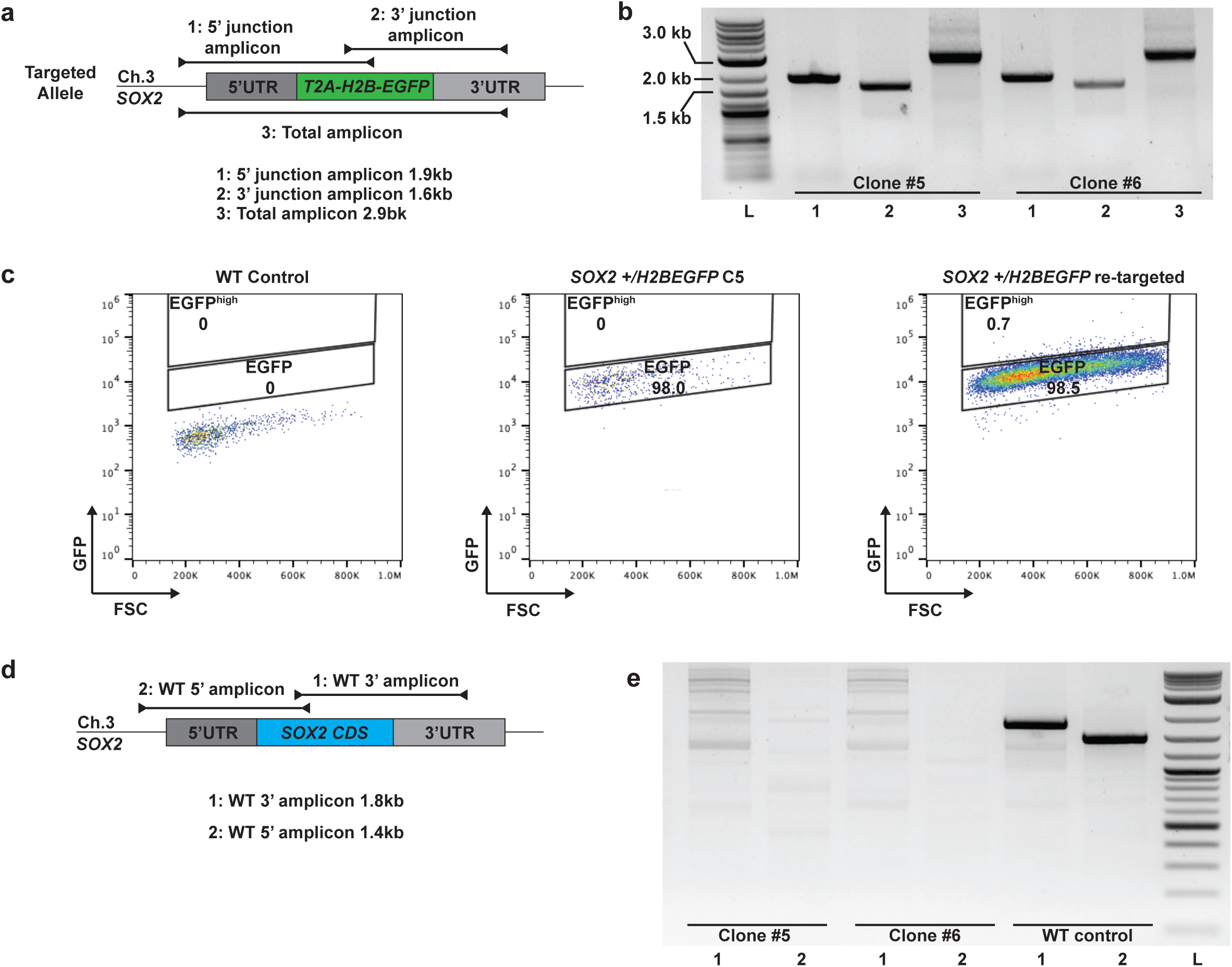
Generation of *SOX2* knockout using organoid Easytag workflow. **(a)** Genotyping strategy for *SOX2 +/H2B-EGFP* heterozygotes. **(b)** Representative gel image showing correctly targeted colonies. **(c)** Representative flow cytometry results showing the EGFP signal shift from WT control to *SOX2 +/H2BEGFP* Clone #5 heterozygote and *SOX2 +/H2BEGFP* Clone #5 retargeted to replace the second copy of the *SOX2 CDS*. **(d)** Genotyping strategy for testing whether the WT *SOX2 CDS* is still present. **(e)** Representative gel image showing correctly targeted colonies (Clone #5 and # 6) with no amplification for WT *SOX2 CDS* in contrast to WT control.

**Supplementary Figure 7.**
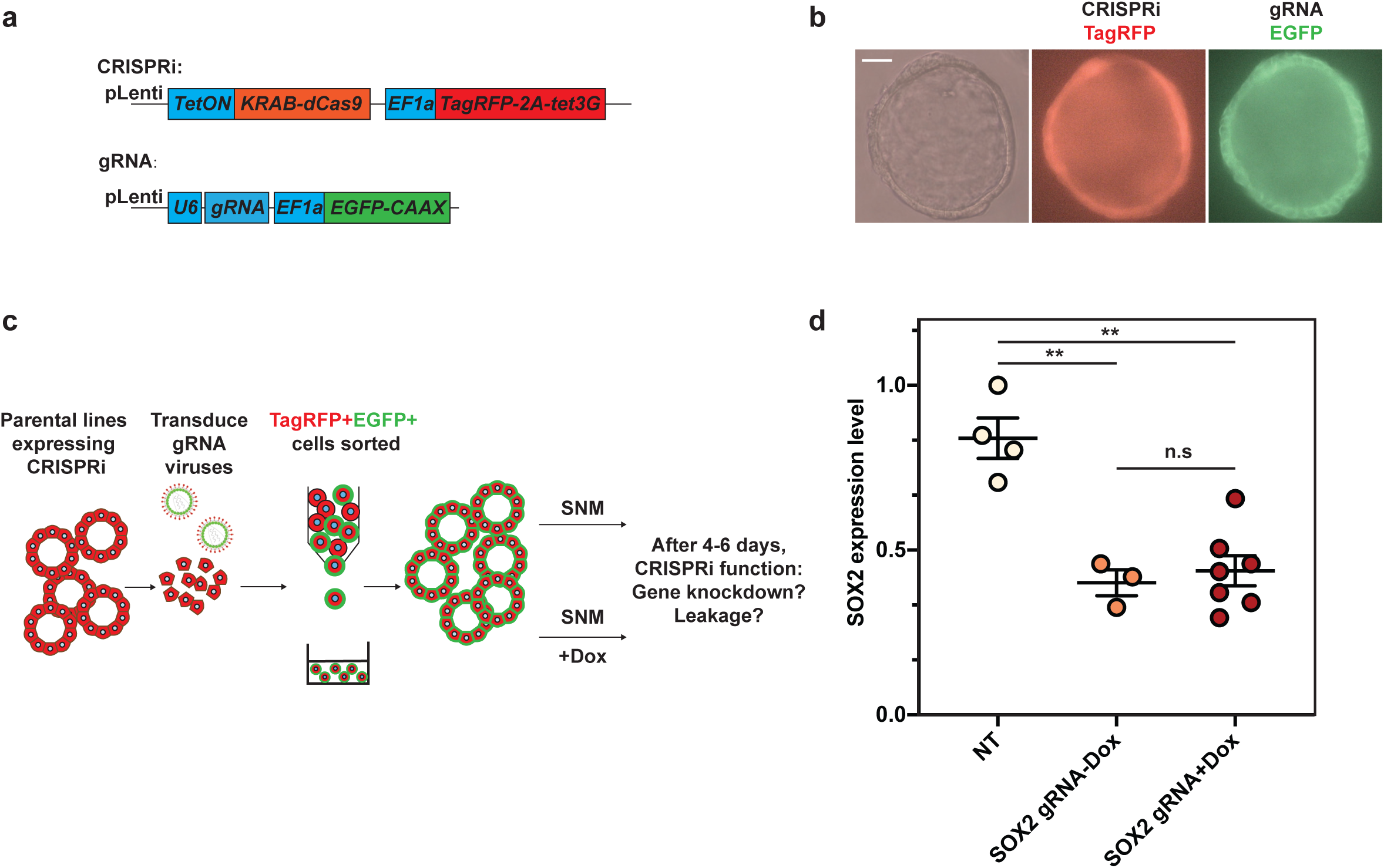
The tetON system alone was not able to tightly control CRISPRi function. **(a)** Schematic of lentiviral design for introducing inducible CRISPRi and gRNA into human foetal lung organoids. **(b)** Representative images showing an organoid colony with both the inducible CRISPRi system and a gRNA. Scale bar denotes 50 µm. **(c)** Workflow for testing the leakiness of the inducible CRISPRi system. A parental organoid line with inducible CRISPRi was introduced via lentiviral transduction, followed by sorting for TagRFP positive cells. Single cells were re-plated (approx. 3000-5000 cells/well of a 24-well plate) and expanded for around 17 days. gRNA lentivirus was then introduced by a second lentiviral transduction event, followed by sorting for TagRFP/EGFP dual positive cells after 3 days. Cells were re-plated again (approx. 2000-3000 cells/well of a 24-well plate) and expanded for another 10 days before treatment with Dox. 4-6 days after Dox treatment, gRNA performance was evaluated. **(d)** *SOX2* can be effectively knocked-down by inducible CRISPRi. However, the knockdown effect cannot be tightly controlled. Organoids with Dox were harvested 4-6 days after the treatment. *SOX2* was effectively knockdown at the mRNA level. However, a similar effect was observed without Dox treatment. Each dot represents an independent experiment. 2 independent organoid lines with 2 different non-targeting control gRNAs and 3 different SOX2 gRNAs were used. SOX2 expression level was normalised to organoids with non-targeting control gRNAs. Error bars are plotted to show mean ± SEM. A two-sided Student’s t-test with non-equal variance was used to compare the means of each group. ** indicates p-value < 0.01. n.s. indicates non-significant.

**Supplementary Figure 8.**
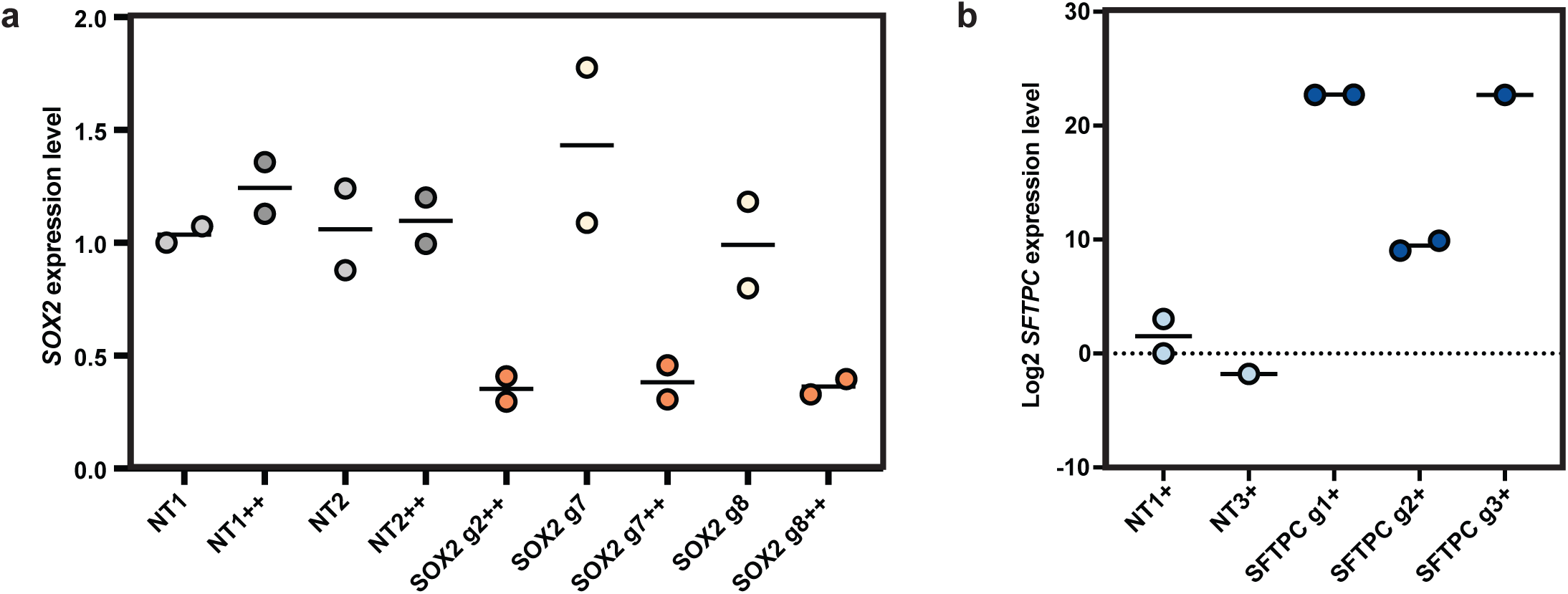
*SOX2* CRISPRi and *SFTPC* CRISPRa results break down. **(a)** qRT-PCR result details for Figure 3 (e). Each dot represents an independent experiment. 2 independent organoid lines were used for each gRNAs. *SOX2* expression levels were normalised to SOX2 expression level in organoids with non-targeting (NT) control gRNA #1. Lines indicate means. **(b)** qRT-PCR result details for Figure 4 (c). Each dot represents an independent experiment. *SFTPC* expression levels were normalised to *SFTPC* expression level in organoids with non-targeting (NT) control gRNA #1. 1 or 2 independent organoid lines were used for each gRNAs. Lines indicate means.

